# Inferring the basis of binaural detection with a modified autoencoder

**DOI:** 10.1101/2021.01.05.425246

**Authors:** Samuel S. Smith, Joseph Sollini, Michael A. Akeroyd

**Affiliations:** Hearing Sciences, Mental Health and Clinical Neurosciences, School of Medicine, University of Nottingham, Nottingham, NG7 2RD, UK

**Keywords:** binaural, hearing, signal detection, representational learning, cross-correlation

## Abstract

The binaural system utilizes interaural timing cues to improve the detection of auditory signals presented in noise. In humans, the binaural mechanisms underlying this phenomenon cannot be directly measured - and hence remain contentious. As an alternative, we trained modified autoencoder networks to mimic human-like behavior in a binaural detection task. The autoencoder architecture emphasizes interpretability and, hence, we “opened it up” to see if it could infer latent mechanisms underlying binaural detection. We found that the optimal network automatically developed artificial neurons with sensitivity to timing cues and with dynamics consistent with a cross-correlation mechanism. These computations were similar to neural dynamics reported in animal models. That these computations emerged to account for human hearing attests to their generality as a solution for binaural signal detection. Methodologically, the study examines the utility of explanatory-driven neural networks and how they may be used to infer mechanisms of audition.

## 1 Introduction

People can detect sounds of interest partially hidden by simultaneously presented background sounds such as noises. The detection of a signal is improved if the time of arrival between the two ears is different to that of the noise (Figure 1, Hirsh, 1948; Licklider, 1948). Detection is typically quantified by measuring the lowest levels at which the signals can be detected (termed “thresholds”), and the improvement here is formally referred to as the “binaural masking level difference” (BMLDs). At low frequencies (Rayleigh, 1907), specialized neural mechanisms utilize interaural time differences (ITDs) less than one ten thousandth of a second to produce BMLDs as large as 15 dB (Hirsh, 1948; Hirsh and Burgeat, 1958). Yet, it is an open question as to what the neurophysiological mechanisms underlying human binaural detection are.

**Figure 1.**
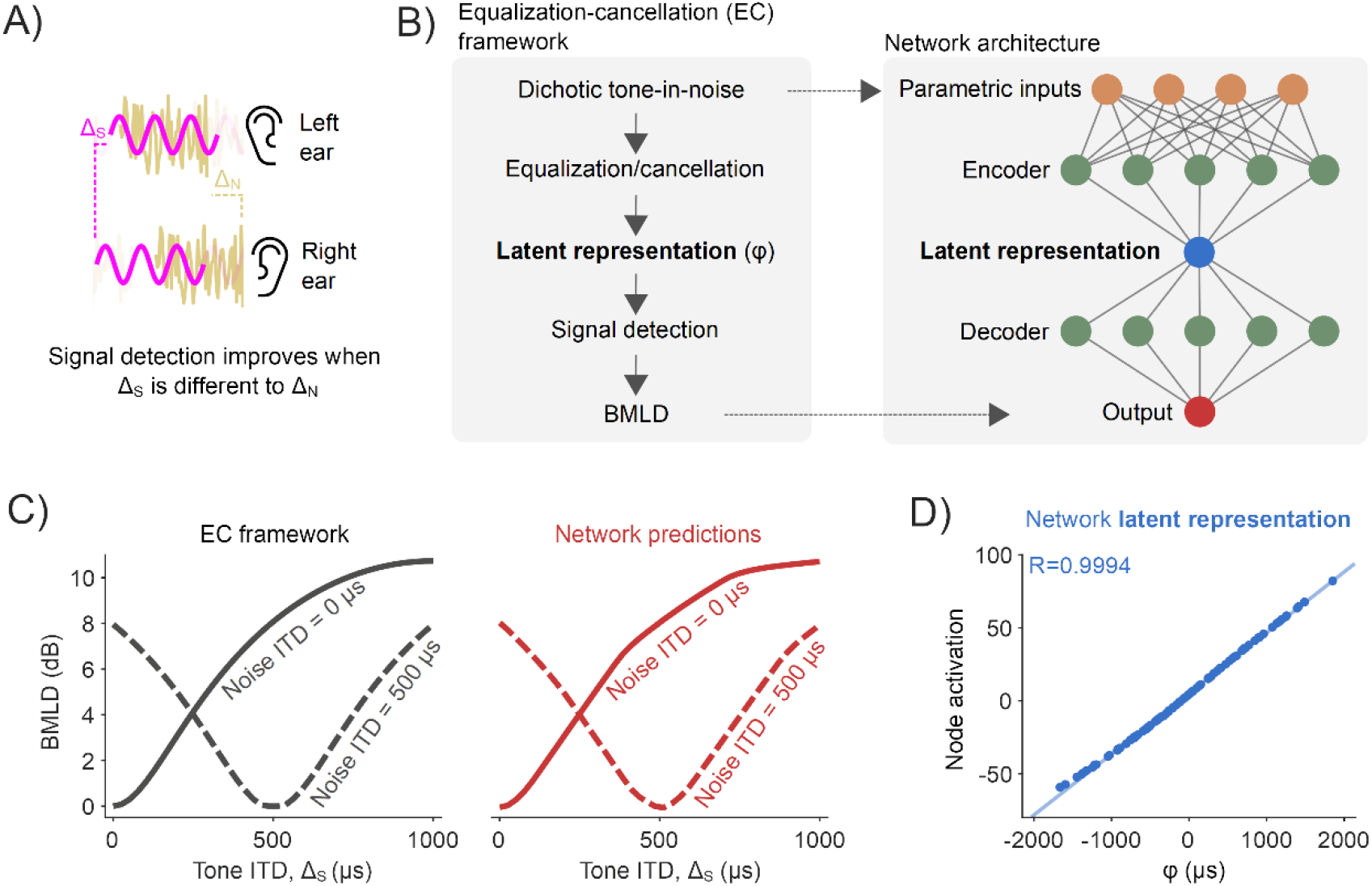
Proof-of-principle: Inferring a latent ITD variable (A)The detection of a signal (pink sine wave) is improved if its interaural disparity is different to that of the noise (yellow noise waveform). (B)A neural network was trained to predict BMLDs, as described by the EC framework (left). The network had a modified autoencoder architecture, in which the central layer acted as an information bottleneck. (C)BMLDs were numerically calculated by the EC framework (black) and estimated by the trained neural network (red), for a 500 Hz pure tone signal and noise at varying ITDs. (D)A node central within the network has activation values entirely consistent with the latent variable as formally defined by the EC framework (φ, the signal’s post-EC ITD).

**Figure 2.**
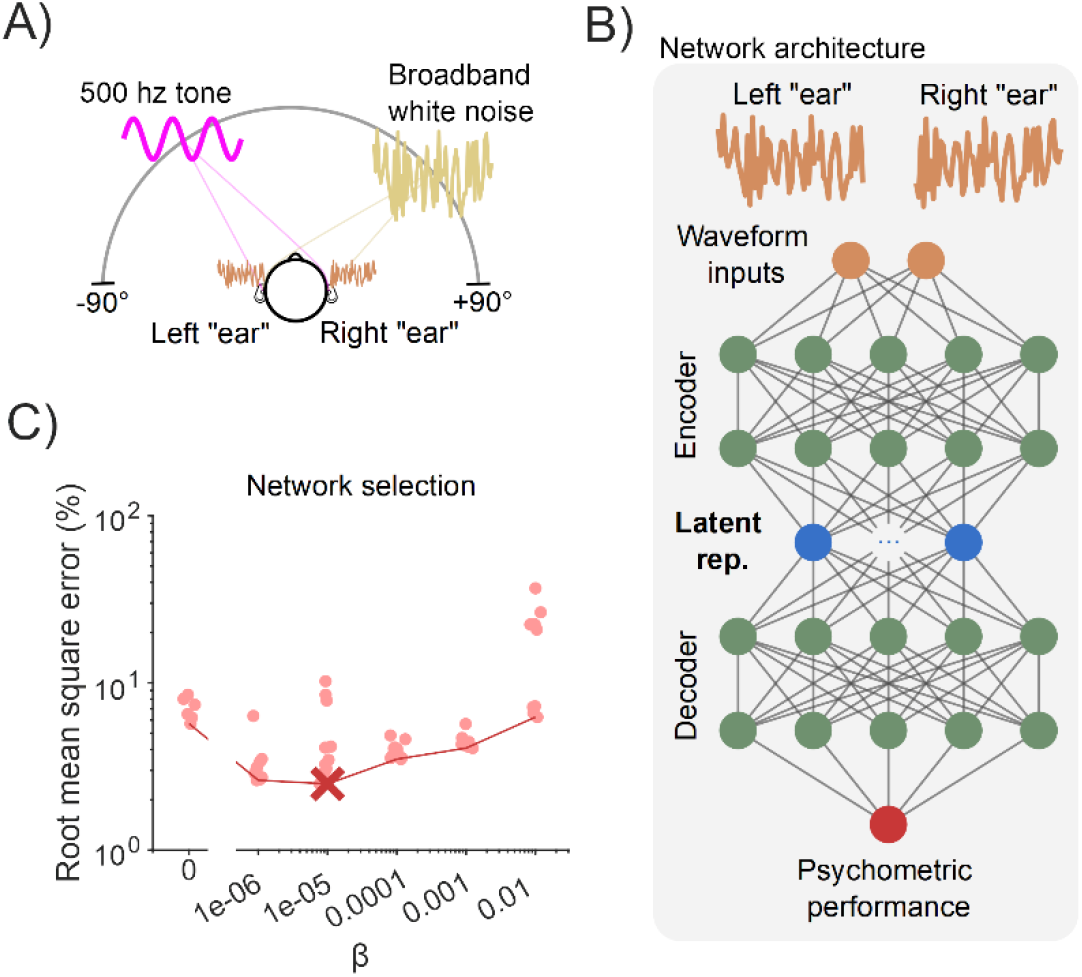
Network training and configuration (A)Data from a simulated frontal field binaural detection task were used to train neural networks to detect a 500 Hz pure tone (pink sine wave) in broadband noise (yellow noise waveform). Locations of the tone and noise were chosen at random on each trial and were equally likely to come from each azimuthal location. (B)The modified autoencoder network received left/right “ear” waveforms as inputs, and had 5 hidden layers, with the central layer containing 10 nodes - constrained by the parameter β in their information transmission. (C)Error for 60 networks (10 for each value of β, see Methods) tested on a held-out validation dataset. The red cross marks the optimally performing network, and the red line bounds the networks with minimum error for each value of β.

For example, midbrain and cortical recordings in non-human species lend support to a cross-correlation treatment of auditory signals across the ears (Palmer and Shackleton, 2002; Lane and Delgutte, 2005; Gilbert et al., 2015). Whereas, human behaviour appears to be equally well, if not better, described by a noise cancellation scheme (Durlach, 1963; Breebaart et al., 2001; Culling, 2007). Discrepancies between frameworks have not been resolved with human imaging data (Sasaki et al., 2005; Wack et al., 2012, 2014; Fowler, 2017), for which resolution and response variability are key limitations. As the neural activity in brain regions underlying binaural detection cannot be directly recorded from in humans, we considered alternative methods of scrutiny.

The human-like “behavior” achievable with deep neural networks, combined with their unpremeditated network of computations, have seen them advocated as a new generation of model organisms (Scholte, 2018). These models are resource efficient, relatively easy to record from and perturb activity in, and are not limited by species-specific ecology. In principle, if a network can be built that corresponds with human behavior, then knowing how that network works might give insight into the human mechanisms. This is effectively treating a neural network as a model organism (Scholte, 2018), and is a promising approach in bridging together neural and behavioral data. Yet, to date, the inner workings of neural networks configured to handle binaural audio have received limited consideration (Adavanne et al., 2018; Vecchiotti et al., 2019; Francl and McDermott, 2022), and almost exclusively in the context of binaural localization rather than detection. One potential stumbling block when interrogating the inner-workings of neural network analogues is their black-box nature. However, network architectures that put mechanistic interpretability at their forefront (such as a modified autoencoders that have shown promise in the field of physics; Higgins et al., 2017; Iten et al., 2018) could help overcome this.

Here, we trained artificial neural networks to imitate the phenomena of binaural signal detection under human-like behavioral constraints, then interrogated their inner workings to discover how they worked. We discovered that not only did the networks learn to make predictions similar to behaviour in humans, but the optimal network was found to have striking similarities with a cross-correlation mechanism similar to animal models (McAlpine et al., 1996; Lane and Delgutte, 2005; Asadollahi et al., 2010; Gilbert et al., 2015). Our key insight -- that these computations emerged to account for human hearing -- attests to their generality as a solution for binaural signal detection and demonstrates the power of representational-learning methods.

## 2 Results

### 2.1 Proof-of-principle: Inferring a latent ITD variable

In this study, we utilized artificial neural networks as a tool to infer key computational variables underlying binaural detection in humans. Such an approach has proven successful in the field of physics (Iten et al., 2018). For example, in the case of predicting the movement of a pendulum, networks have correctly inferred an influential role of variables such as spring constant and damping factor. To demonstrate the feasibility of this methodology in the context of binaural hearing, we trained a network on a reduced example. We wanted to verify that, in the process of predicting the dynamics of a fully defined system, the network would infer the same latent variable within that system. Namely, we trained networks to mimic a system of equations derived under the “equalization-cancellation” (EC) framework (Durlach, 1963), equations effective at reproducing human binaural psychophysics (Durlach, 1963; Klein and Hartmann, 1981; Breebaart et al., 2001; Hartmann and McMillon, 2001; Culling, 2007; Wan et al., 2010). The framework proposes that the interaural configuration of the masking noise is “equalized” (applies an internal time delay to the waveform from one ear to compensate for, or equalize for, the external temporal disparity compared to the waveform from the other ear) and “cancelled” (subtracts the equalized waveforms from one another), resulting in a more detectable signal. These EC operations give rise to a latent representation that can be captured by the variable φ, describing the ITD of the post-EC signal (Figure 1B, left). In the EC framework, this variable is used in the equations to predict the consequent improvement in signal detection, i.e., BMLDs. We were interested as to whether a neural network would automatically infer the latent variable φ in the process of predicting BMLDs as described under the EC system of equations.

We trained a modified autoencoder neural network to predict the *binaural* improvement in signal detection (i.e. BMLDs) based upon four parameters describing the *monaural* arrival times of a 500 Hz signal and broadband noise at each ear, taken from the input to the EC equations (Figure 1B). Following training, we found that the neural network was able to generalize its performance to predict BMLDs for parametric combinations for which it had not been trained (root mean square error of 0.075 dB on the validation dataset). The network correctly predicted larger BMLDs when the signal had a non-zero ITD and the masking noise did not, and vice versa (Figure 1C). Interrogating the computations latent within the network provides insight into how it operates. Because the network utilized a modified autoencoder architecture, its inputs are “encoded” into a simpler representation, the latent representation, by passing information through a bottleneck at the center of the network (Figure 1B, right). When we look at the bottleneck node’s activation values (its numerical readout), we see that its activation entirely corresponds to the latent variable in the EC framework, φ (Figure 1D; Pearson’s R = 0.9994, p<0.001). In summary, within this fully defined system, the network was able to infer the appropriate latent variable in accounting for BMLD dynamics, and therefore confirm out premise.

### 2.2 Modified autoencoder accounted for binaural detection psychophysics

In Section 2.1 we provided the network with four parameters quantifying a signal in a noise, whereas the human auditory system would be presented *waveforms* of a signal combined with masking noise – each in unknown quantities. How these waveforms are processed as to confer a binaural advantage is an open question, nor does the EC framework make any explicit proposal about how equalization parameters would be derived from said waveforms (Durlach, 1972; Wan et al., 2010). Additionally, humans display a graded psychometric performance as signal level is varied, from inability to full detection, for which detection thresholds only offer a single-value snapshot at one chosen performance level. We advanced the network/training paradigm described in Section 2.1 to incorporate these aspects of binaural detection – namely, left/right “ear” waveform inputs and graded psychometric-function outputs. We also generalized the training data to represent signals coming from random azimuthal locations in the frontal horizontal plane, restricting the range of incorporated ITDs to within an approximate human physiological range (±655 μs; Figure 3a). To generate BMLD estimates, we retained the set of equations used in Section 2.1 (using parameters from which waveforms were constructed), as they represent good fits to human binaural psychophysics and augment the availability of training data. To account for the increased complexity, the autoencoder was modified to have two layers of artificial neurons at the “encoder” and “decoder” stages and allowed for multiple (10) nodes in the central latent representation part of the network (Figure 3b). We ran 60 separate networks, each trained on the same data, but ran with varying constraints as to how independently each latent node represented information, set by parameter β that balanced network regularization versus network interpretation. When validating the predictive accuracy of the networks, we found that networks with an intermediate value of β best accounted for a held-out set of data (Figure 3C), showing that some constraints on information encoding were better than none.

**Figure 3.**
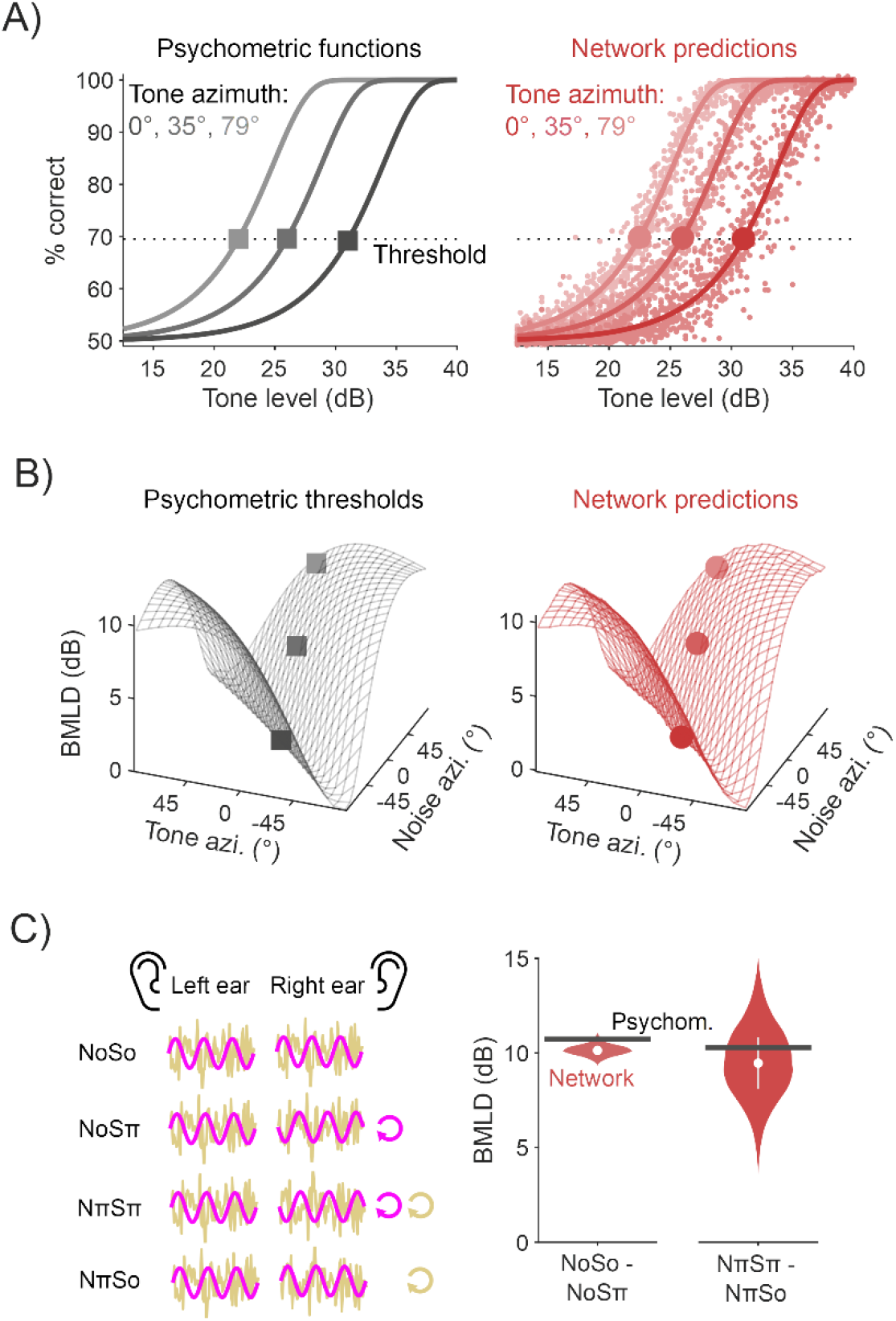
Modified autoencoder accounted for binaural detection psychophysics (A)Psychometric functions quantifying tone detection as a function of tone level masked by a 60 dB Gaussian noise (left, black). These functions are drawn for tones presented from three azimuths, relative to a noise presented directly in front. The optimal neural network was able to approximate these psychometric functions (red, right), from which detection thresholds (corresponding to a d-prime of 1) and BMLDs could be calculated. (B)Psychophysical estimates (left, black) of human BMLDs for 500 Hz a tone presented in noise, each with ITDs mapped from differing azimuths. Alongside are the optimal network’s predictions (right, red). Markers representing thresholds as defined in 3A are overlaid. (C)BMLDs were derived for experimental stimulus configurations: NoSo/NoSπ, NπSπ/NπSo (Note: π, for a 500 Hz signal, is beyond the range of ITDs used during training).

The optimal network had a root mean square error of 2.5% for the validation dataset. We found this network was able to replicate the psychometric functions for the improvement in signal detection as the presented tone increased in level (Figure 3A) in a 60 dB broadband noise. From these data, we were able to regress functions from which to derive detection thresholds (defined as a performance level of d-prime of 1) and, in turn, calculate BMLDs. We found that the network BMLDs decreased as the difference between tone ITD and noise ITD increased (Figure 3B). For example, in diotic noise (noise ITD = 0) with a tone placed at the far left, detection thresholds were significantly enhanced by 9 dB (two-sided unpaired t-test, p<0.001).

To allow comparative assessment of the neural network and previously published work, we also tested the network on stimulus configurations typically employed in the laboratory to study binaural detection. These include tones and noise in extreme configurations, either in-phase or completely out-of-phase across the ears. In the literature these stimuli are denoted as NoSo, NoSπ, NπSπ, and NπSo, where N refers to the noise, S the pure tone signal, with the subscripts denoting interaural phase difference (IPD) in radians. These stimuli have ITDs that are frequency dependent and can be greater than the range permitted by head width. For example, a 500 Hz pure tone with an IPD of π corresponds to an ITD of 1000 μs, whereas the typical value for the largest ITD due to a head is 655 μs (Woodworth et al., 1954). As our networks were trained on ITDs within the head’s range, this meant the network had no prior exposure to this magnitude of ITD and so it was unclear how it would function over this range. We found that when the noise signal had zero IPD, the BMLD for the corresponding homophasic (NoSo) and antiphasic (NoSπ) tone conditions was 10.1 dB (two-sided unpaired t-test, p<0.001). Comparatively, when instead the noise signal was interaurally out-of-phase, the predicted BMLD for the corresponding homophasic (NπSπ) and antiphasic (NπSo) stimuli was 9.5 dB (two-sided unpaired t-test, p<0.001). These BMLDs are similar to those measured in people (Durlach and Colburn, 1978) and with estimates from the psychophysical equations (10.7 dB and 10.3 dB respectively; Figure 3C).

### 2.3 Latent representations imitate neural signature of population-level cortical activity

Given the network’s human-like performance over a range of trained, and untrained, conditions (see Section 2.1), we next investigated *how* the model achieved this “behavior”. To do this we, first, looked to the network’s latent representations and considered them relative to known binaural phenomena. Namely, animal neural data have shown that the stimulus conditions depicted in Figure 3C (NoSo/NoSπ and NπSπ/NπSo) hint at a unique signature of binaural detection processing (Gilbert et al., 2015). Despite both NoSo/NoSπ and NπSπ/NπSo conferring similar benefits in human binaural detection (Figure 3C), different neural dynamics appear to be associated with these stimulus conditions. In guinea pig cortical recordings, population spike counts dropped amongst an No signal as a 500 Hz tone went from So to Sπ (Figure 4A). Conversely, amongst an Nπ signal, as a pure tone transitioned from Sπ to So, population spike counts *increased*. The neural dynamics contrast, yet the behavioral dynamics do not. However, the neural and behavioral measures emanate from different species and hence their contrast may be a consequence of this rather than mechanistic in nature.

**Figure 4.**
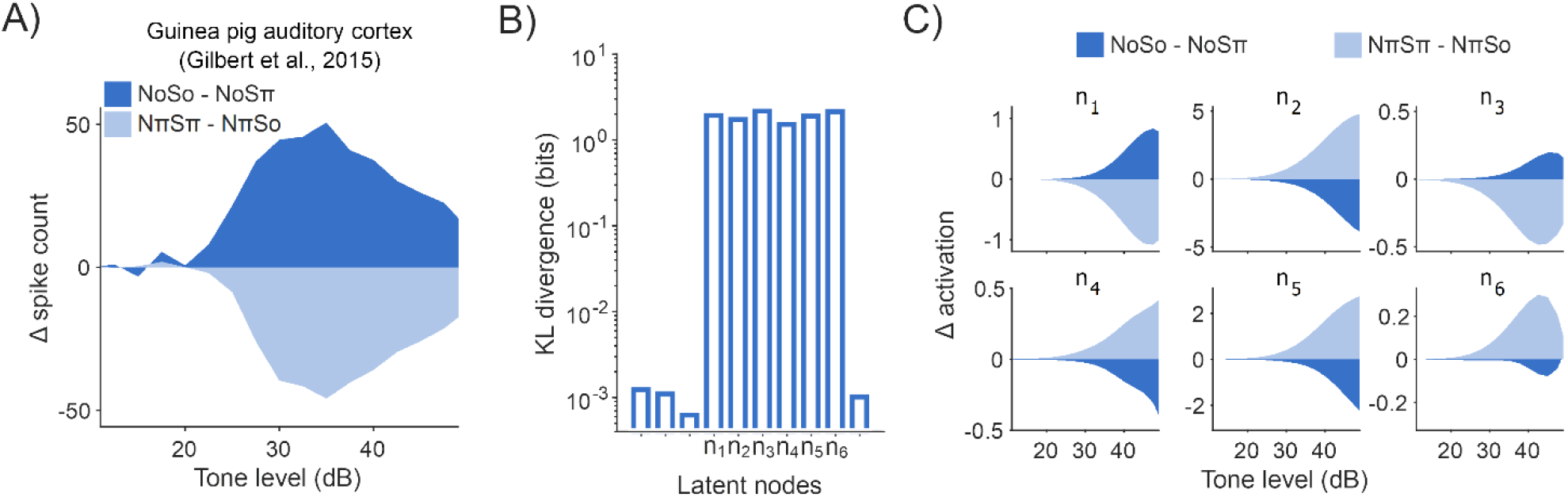
Latent representations imitate neural signature of population-level cortical activity (A)Population masked rate-level functions recorded from guinea pig auditory cortex (Gilbert et al., 2015) in response to changes in experimental binaural stimuli NoSo/NoSπ (dark blue) and NπSπ/NπSo (light blue). (B)Kullback–Leibler (KL) divergence (Kullback and Leibler, 1951) between each individual node and a unit Gaussian. Unlabeled nodes along the x-axis were deemed to be suppressed during training. (C)Rate-level functions for the operational latent nodes in the optimal network, comparable to Figure 4A.

We therefore examined how NoSo/NoSπ and NπSπ/NπSo stimulus conditions were represented in the central latent layer of our network. Following training, six of the ten central nodes were deemed operational, in the sense that they had nontrivial output values, whereas the remaining four had adapted to produce negligible outputs to comply with constraints on information transmission (Figure 4B). We found that these nodes exhibited opposing dynamics in response to the two pairs of homophasic/antiphasic stimuli (Figure 4C), although, the directionality of these opposing dynamics varied across the six nodes (we believe that this is a consequence of the nodes being able to take any real number, and hence this directionality can be ignored). On average, the change in activation for NoSo/NoSπ was opposite to NπSπ/NπSo for all six operational nodes (a 2^-6^ = 0.016 chance). Although mean differences were significant (two-sided unpaired t-tests for tone-level of 35 dB, p<0.001 for all), trial-to-trial latent values were noisy and overlapping (two-sample K-S test between NoSo/NoSπ and NπSπ/NπSon conditions, for a tone level of 35 dB, D ranged from .076 (n4) to .43 (n2), p<0.001 for all), to be expected given the input waveforms were dominated by Gaussian noise. Some of this variance was due to the partial representation of non-binaural stimulus properties (e.g. tone phase) that had not been adequately disregarded early in the network. Some of this variance could be regressed out based upon the activity of other latent nodes (Figure 4SA). With such co-variation accounted for, we saw a further enhanced contrast for the NoSo/NoSπ and NπSπ/NπSo stimulus conditions, markedly at threshold levels (Figure 4SB,C). In summary, given that the network predicted similar magnitudes of BMLDs for NoSo/NoSπ and NπSπ/NπSo, *and* broadly captured opposing latent dynamics for these stimulus conditions, we conclude that the network imitated this key signature of binaural detection and reconciles the signature within a single model.

**Figure S4.**
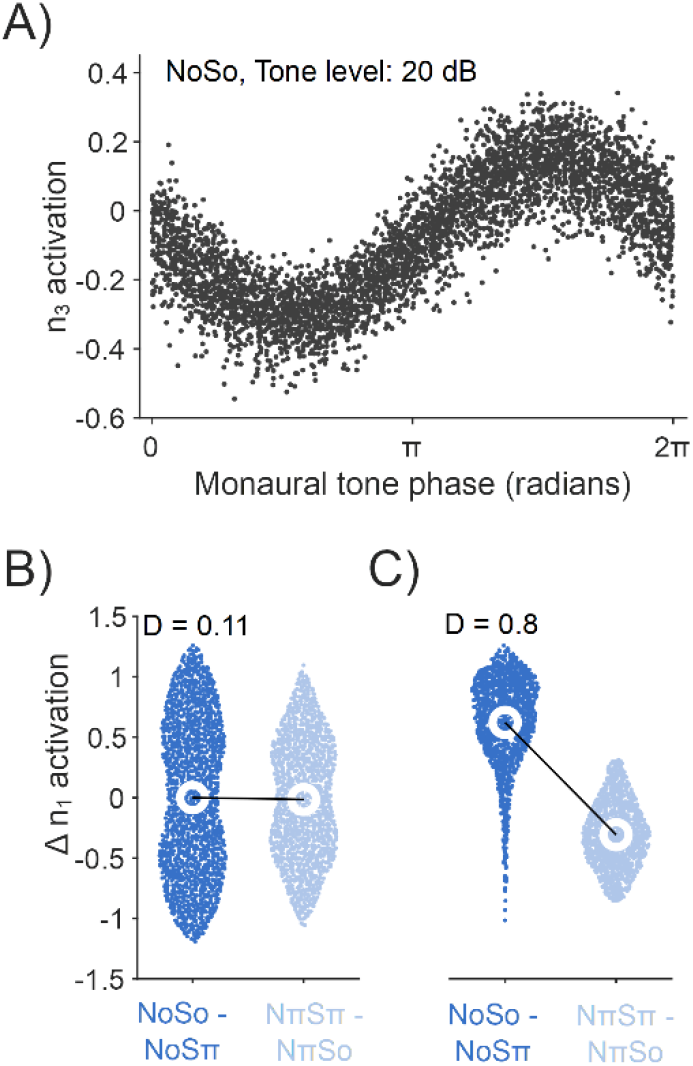
Supplementary. (A)Some latent nodes orthogonally represented stimulus-properties. For example, n3 sinusoidally varied in activation value as a function of monaural tone phase. Shown for NoSo with tone level at 20 dB. (B)Near threshold (tone level of 20 dB), the distribution of values when comparing the change in n1 activation between NoSo/NoSπ (dark blue, left) and NπSπ/NπSo (light blue, right) are considerably overlapping. Two-sample KS test statistic, D, is 0.11, p<0.001. (C)When the co-variate captured by n3 is controlled for (e.g., looking at when n_3_ < 0, i.e., monaural tone phase between 0 and π) the distinction between the conditions is clearer. Two-sample KS test statistic, D, is 0.8, p<0.001.

### 2.4 Encoder network dynamics matched those of a cross-correlator

To further understand the encoder network that lies between the waveform inputs and the central latent representations, we examined ITD tuning within the network. To determine this, we recorded noise delay functions in artificial neurons (Figure 5A), i.e. their activation values in response to noises presented with varying ITDs, as these are typically measured in animal physiology studies (Joris and Yin, 2007). Tuning was quantified by regressing a Gabor function (Lane and Delgutte, 2005), the combination of a cosine windowed by a Gaussian (overlaid in Figure 5A). For nodes in the encoder’s first layer, we observed significant ITD tuning in 63 out of 100 nodes (Figure 5B). By the encoder’s second layer, significant ITD tuning had emerged in all 100 nodes. Estimates of each node’s best ITD (i.e., the ITD that gives the maximum activation) were derived from Gabor fits (to account for nodes that were cyclical in their noise delay responses, best ITD was attributed to the most central tuning peak). In both the first and second layers of the encoder network, we observed a wide distribution of best ITDs, both within the simulated head-range, and beyond it (Figure 5C).

**Figure 5.**
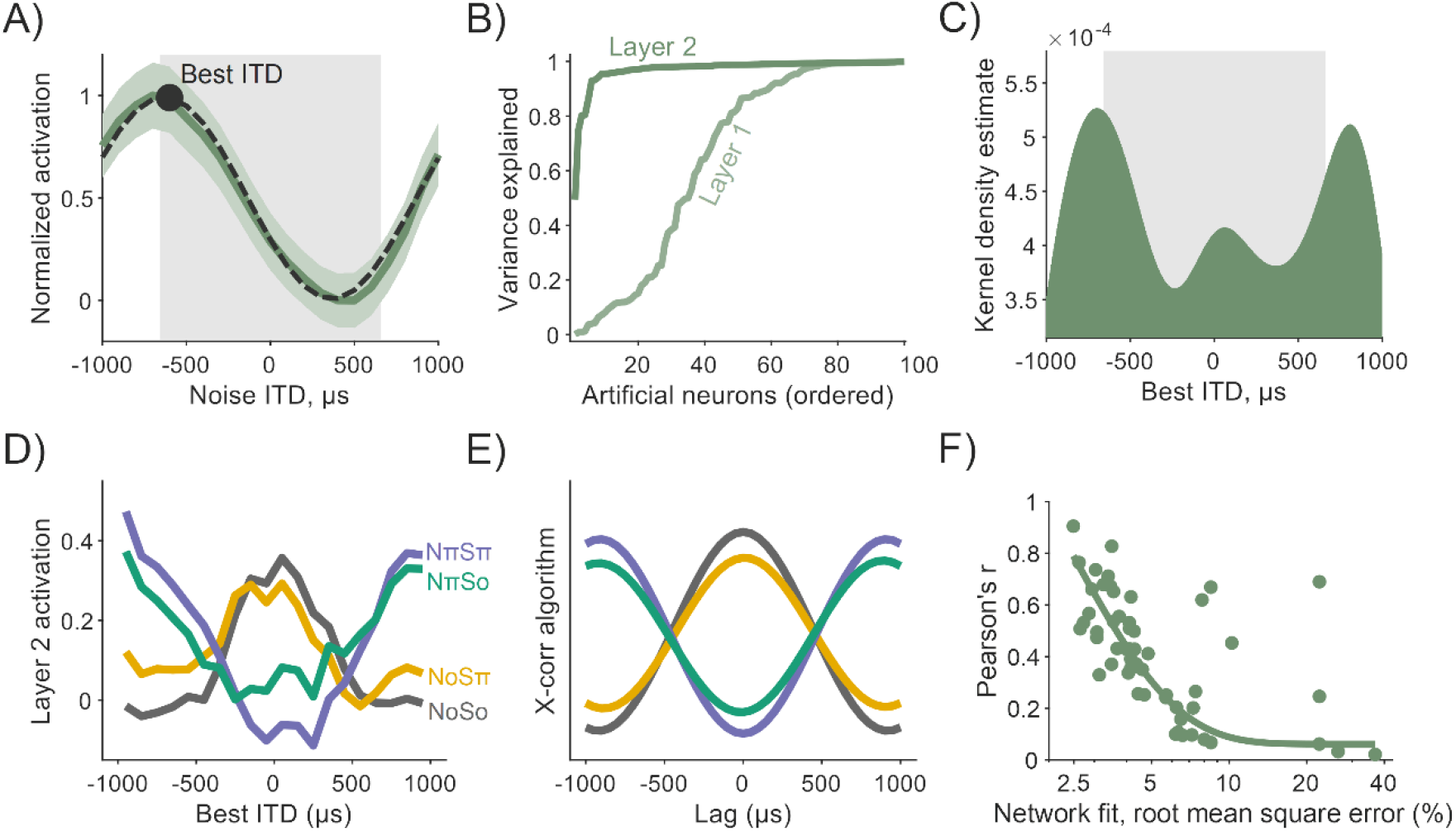
Encoder network dynamics matched those of a cross-correlator (A)ITD tuning emerged as a property of nodes within the early encoder layer of the network. Node activations of an example artificial neuron are shown to vary as a function of noise ITD (dark green). Tuning was characterized by Gabor functions (black, dashed) with peaks defined as an artificial neuron’s best ITD (black circle). The gray box underlays represent the ITD-limit for our simulation. (B)The proportion of variance explained (R^2^) by Gabor fits, although high in Layer 1 (light green) of the encoder, was widespread by Layer 2 (darker green). (C)Best ITD distribution for artificial neurons in Layer 2, characterized by a kernel density estimate (bandwidth of 200 μs). Again, the gray box underlay represents the ITD-limit for our simulation. (D)Activation values of Layer 2 artificial neurons for binaural detection stimuli: NoSo, NoSπ, NπSπ, NπSo (color-coded). Smoothed with a 600 μs moving average window. (E)The profiles in 5D were similar to a simple cross-correlation (X-corr) algorithm. (F)The better a network predicted psychophysical data (x-axis), the more similar its encoder network to a cross-correlator.

One framework that is commensurate both with the opponent dynamics described in the latent nodes (Section 2.3) and found in animal models, is that of a binaural cross-correlator mechanism (McAlpine et al., 1996; Lane and Delgutte, 2005; Asadollahi et al., 2010; Gilbert et al., 2015). The concept is predicated on the existence of coincidence detectors that encode temporally offset signals. To deduce whether the network had automatically learnt to operate like a cross-correlator, we measured artificial neuron activations in responses to the extreme tone-in-noise conditions: NoSo, NoSπ, NπSπ, and NπSo (Figure 5D). When a signal was presented amongst an in-phase noise (No), responses were largest for artificial neurons with best ITDs near 0 μs and decreased as best ITDs were increasingly non-zero. Conversely, amongst an out-of-phase noise (Nπ), responses were lowest for nodes with best ITDs near 0 μs and increased as best ITDs deviated away from this. The effects of tone phase on node dynamics were more subtle, although these dynamics were also in accordance with a nodes’ tuning properties. Nodes tuned to smaller ITDs responded most to in-phase tones (So) and least to out-of-phase tones (Sπ), and vice-versa for nodes tuned to larger ITDs. These dynamics are consistent with a cross-correlation model.

Computationally, a binaural cross-product can be calculated by summing the point-by-point product of two temporally offset signals. Comparative outputs from a simple binaural cross-correlation algorithm (namely for signals in noise passed through narrow-band filters centered at 500 Hz) are shown in Figure 3E. We saw a significant correlation between the network and the cross-correlation calculation (with local averaging: Pearson’s r=0.91, p≪0.001; without: Pearson’s r=0.36, p≪0.001). When looking across all 60 of the networks that we trained, we found that the more a network made predictions similar to the psychophysical data, the more similar its encoder network was to a cross-correlator.

## 3 Discussion

Binaural detection of a signal masked by noise is a well-standardized laboratory measurement that underpins important theories of auditory processing. However, the underlying mechanisms involved remain uncertain. Here, we used for the first time representational learning methods to infer potential mechanisms underlying human-like binaural detection. We found that our neural networks were able to successfully utilize interaural discrepancies across dichotic signal-in-noise waveforms to predict human-like binaural detection behavior. Notably, similarities with a binaural cross-correlator and animal neural dynamics were emergent within the network. These dynamics were not hard-coded into the network and highlight their importance in the context of signal detection, not just the more commonly referenced function of sound localization (Joris and Yin, 2007). These findings promote the understanding of how neural networks work as an effective tool for investigating the basis of auditory processing.

### 3.1 The basis of binaural detection

In our methodology, we utilized a set of equations originally derived from the assumptions of the EC framework (Durlach, 1972), treating them as accurate phenomenological fits to human binaural psychophysical data. This is the case, and was in part the motivation, for the experimental parameters investigated in this study (i.e. a 500 Hz tone, ITDs but not ILDs (Wan et al., 2010)). However, our discoveries overall support a different process for interaural detection, namely cross-correlation. The distinction is important because Domnitz and Colburn (1976) provided statistical evidence that, under certain assumptions, models based on temporal or phase differences (as the EC framework is) provide similar predictions of tone-in-noise detection to interaural correlation-based models. They thus concluded that comparing binaural detection predictions made by both classes of models is insufficient to disentangle underlying mechanisms. To circumvent this, we inverted the conventional forward-approach to modelling, and instead effectively reverse engineered our models. We discovered that our models developed a cross-correlation mechanism to reproduce psychophysical data. We also observed that latent nodes broadly reproduced opposing dynamics to NoSo/NoSπ and NπSπ/NπSo, matching what has been found to exist in animal models. In contrast, one would expect that an EC-like noise cancellation scheme would operate similarly for both NoSo/NoSπ and NπSπ/NπSo stimulus conditions, and hence would not exhibit these opposing dynamics. Further, we found that additional mechanisms that utilize *a priori* knowledge of the masker, as has been proposed for some EC models (Hawley et al., 2004), are not necessary to account for binaural detection behavior. Taken-together, one interpretation of our findings is that binaural cross-correlation represents an implementation of the “computational goal” of the system (Marr and Poggio, 1976), where the computational goal may coincide with the motivations underlying the EC framework (i.e., to increase the signal-to-noise ratio by eliminating the noise).

Despite the occurrence of the aforementioned network dynamics, we also observed dynamics that were less tangible. For example, we observed instances in which latent nodes partially represented seemingly irrelevant stimulus properties, e.g., monaural phase. As opposed to the encoder network filtering out these stimulus properties, the network opted to separately account for this co-variation and account for it at a later stage. This is likely a consequence of the modified autoencoder architecture’s preference for capturing separate latent variables in separate nodes (Higgins et al., 2017; Iten et al., 2018), potentially augmented by an over resourced “decoder” network. In addition to these divergent latent dynamics, for some extreme stimulus configurations we observed some slight discrepancies in predicted and ground truth detection thresholds, although we stress that relative differences (i.e., BMLDs) were accurately predicted. One alternative is that networks could be trained to perform binaural detection for stimuli configurations with ITDs beyond the limits imposed by a typical head size, to mimic distributions that incorporate statistics of natural sounds that can be more numerous and overlapping (Młynarski and Jost, 2014).

### 3.2 Neural network analogues of auditory processing

Understanding of binaural detection in humans has been mired due to ambiguity regarding whether animal neurophysiology data satisfactorily accounts for human psychophysics. Treating deep neural networks as a model organism (Scholte, 2018) appears to be a promising approach in bridging together neural and behavioral data. Recent neural network studies have described correlates with broad organizational principles in the auditory system (Kell et al., 2018; Koumura et al., 2019; Khatami and Escabí, 2020) and asked questions of “why” a neural system operates in a particular way. Here, we focused on the question of “how” a system operates, for the well characterized binaural phenomena of improved detection of 500-Hz tone in noise. Despite the notable computational similarities between our trained networks and neural observations, comparisons between artificial neural networks and neural biology come accompanied by an asterisk. The network was not constructed with the goal of accurately mimicking neuronal biophysics or hierarchical complexity, but instead a trade-off was made in which a modified autoencoder architecture (Iten et al., 2018) was applied to facilitate interpretation and optimization. Future work, scaling this modelling approach, for example to examine interaural level differences or across-frequency integration, would likely be insightful. However, any impact on interpretability should be weighed (even in this, arguably simplified, context the network dynamics were non-trivial), and such models are first contingent on the generation of suitably large psychophysical datasets.

### 3.3 Conclusion

In conclusion, our results demonstrate for the first time that an artificial neural network, utilizing a modified autoencoder architecture, can discover key computations underlying binaural hearing. Key latent activity within the model corroborates observations made in animal physiology and speak to their generality as a solution to binaural detection. The work demonstrates the potential for representational learning methods to help bridge the gap between neurophysiology and psychophysics.

## 4 Materials and methods

### 4.1 Binaural detection rates and thresholds

The framework of equalization and cancellation (Durlach, 1972) has wide human psychophysical support, successful in predicting binaural masking level difference (BMLD) data (Durlach, 1972), binaural pitch phenomena (Durlach, 1972; Klein and Hartmann, 1981; Hartmann and McMillon, 2001), underpinning other models of binaural hearing (Breebaart et al., 2001), and proven psychophysically favorable relative to other prominent theories (Culling, 2007). It is therefore a suitable “standard theory” to build our work on. Numerical predictions of BMLDs in decibels were calculated from phenomenological equations derived from this framework (Durlach, 1972; Wan et al., 2010):

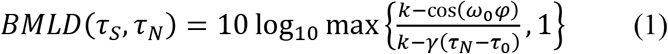

where *τ_S_* and *τ_N_* are the interaural time lags of the signal and noise, *ω*_0_ is the angular frequency of the pure tone signal, 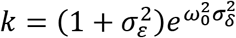 where 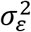 and 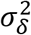 are jitter (internal noise) parameters, γ is the normalized envelope of the autocorrelation of the narrow-band noise output of a filter centered at the target tone frequency, and *τ*_0_ is an optimal time equalization parameter. The parameter *φ* = *τ_S_* – *τ_N_* represents the difference in interaural time of the tone and noise signals; in Section 2.1, we examined whether a neural network could discover this parameter. The values of the other parameters were chosen according to Durlach’s (1972) original formulation in which the model was fit to human data, e.g. *γ* assumes a filter with triangular gain characteristics.

Psychometric functions were derived under the assumptions that (i) BMLDs are relative to a nominal diotic detection threshold of 31 dB, and (ii) detection thresholds were for a d’ of 1 in a yes-no experiment (Ingleby, 1967; Egan et al., 1969):

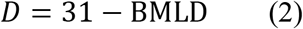

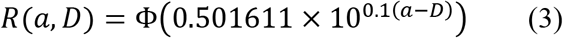

where BMLD is from Equation 1, *a* is pure tone amplitude, *D* the detection threshold in dB, and Φ is the inverse cumulative normal distribution.

### 4.2 Modified autoencoder network

We ran autoencoder-based deep neural networks (Higgins et al., 2017; Iten et al., 2018). Networks took input values that were passed through exponential linear unit (ELU) layer(s), referred to as the “encoder” portion of the network. This was followed by a single layer with latent Gaussian node(s) (≥ the number of parameters varied in the generation of training stimuli) with minimal uncorrelated representations, constrained by a parameter *β* which balances network regularization versus network interpretation. This was followed by further ELU layer(s), referred to as the “decoder” portion of the network. All layers were fully connected and feed-forward. The Adam optimization algorithm (Kingma and Ba, 2014) was used to minimize the cost function:

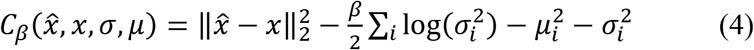

where 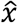 and *x* are predicted and ground truth outputs respectively (subscript 2 is the L2 norm, superscript 2 is squaring), *σ* and *μ* are the standard deviation and mean of latent Gaussian nodes respectively, and the *i* subscripts reference separate latent nodes. Architecture meta-parameters were influenced by those described in Iten et al. (2018). Networks weights/biases were randomly initialized. Batch size (number of training instances employed in each iterative update of network parameters) was set to 256. The learning rate (training hyperparameter) was set to 5×10^-4^ for 1000 epochs (total passes of entire training dataset).

### 4.3 Parameter-based network

The “proof-of-principle” network (see Section 2.1) took 4 parametric inputs representative of arrival times of a 500 Hz pure tone and broadband noise at each ear: arrival times of the pure tone at the two ears and the arrival times of the noise at the two ears. The network was trained to predict BMLDs, as specified in Equation 1. The network was trained and validated (95%/5% split respectively) on 100 000 instances of monaural tone and noise arrival times, each randomly drawn from between 0 and 2000 μs. The encoder and decoder portions each had one layer with 100 ELU nodes. The latent layer had two nodes (one was suppressed during training) with β set to 10^-5^.

### 4.4 Waveform-based networks

All other neural networks took 800 input values, representative of simulated left “ear” and right “ear” waveforms, each of 400 samples as simulated from a pure tone and noise mapped to different angles in the azimuth. Networks were trained to predict the corresponding detection rates, as specified in Equation 3. Training/validation (95%/5% split respectively) was performed with 1 000 000 instances of a random phase tone in randomly generated white noise. Pure tones had 10 periods, completing one period per 40 samples. Pure tones were treated as 500 Hz for generating estimates in Equation 1. Pure tones were set to levels between 0 and 50 dB. Pure tones were masked by randomly distributed broadband noise (50-5000 Hz, limited by 6th order Butterworth bandpass filter) with an overall level of 60 dB. The tone and noise were gated simultaneously. Tones and noises were simulated with ITDs mapped from two independent angles in the azimuth between -90° (far left) and 90° (far right). ITDs were derived from Woodworth’s equation (Woodworth et al., 1954), assuming a head radius of 0.0875 m.

The encoder and decoder portions of the network each had two 100-neuron ELU layers. The central layer of networks had 10 latent Gaussian nodes. Ten networks were trained for each value of β, namely 0, 10^-6^, 10^-5^, 10^-4^, 10^-3^, and 10^-2^ giving 60 in total. The network with the least root mean square error between predicted detection rates and ground truth for the validation dataset was selected for further analysis. Central nodes were considered operational if the Kullback–Leibler divergence (Kullback and Leibler, 1951) between their individual responses and a unit Gaussian was larger than 0.1 bits.

### 4.5 Network predictions

BMLD predictions were generated by averaging outputs for 10 repeats of a given stimulus configuration (i.e., stimulus ITDs would be fixed whilst other parameters were randomized 10 times). For the waveform-based networks, BMLDs had to be derived based upon detection rates. To determine detection thresholds, the mean of 10 detection rates for tone levels, set between 0 and 50 dB in 2.5 dB steps, were regressed with a psychometric curve (Equation 3; Figure 3A). BMLDs were predicted for (i) random phase tones amongst randomly generated broadband noise with ITDs each mapped from 27 fixed azimuthal locations spaced between ±90°, and (ii) random phase tones amongst randomly generated broadband noise each either in- or out-of-phase (i.e. NoSo, NoSπ, NπSπ, and NπSo).

### 4.6 Artificial neural representations

Artificial neural activation values (= a neuron’s numerical expression) were measured in response to the stimuli configurations in Section 4.5. In addition, activation values were measured as a function of ITD for broadband noise only (50-5000 Hz, 60 dB). ITDs ranged from -2000 μs to 2000 μs in steps of 100 μs. For the parametric-based network, latent activations were measured in response to 100 random stimulus generations. For the waveform-based networks, activations were measured in response to 5000 random stimulus generations.

### 4.7 ITD tuning

ITD tuning was quantified by fitting a Gabor function (Lane and Delgutte, 2005) to noise delay responses. The parametric expression for a Gabor function is:

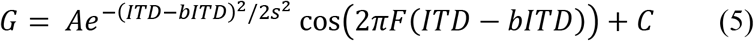

in which we characterized a nodes’ best ITD as the parameter *bITD, F* is the tuning curve frequency, *A* is a scaling factor (constrained to be positive), *C* is a constant offset, and s is a decay constant. These parameters were fit with the non-linear least squares algorithm curve_fit (a SciPy function; Virtanen et al., 2020). An F-test was used to assess whether a Gabor function was a significantly better fit to noise delay responses than a linear function of ITD.

### 4.8 Binaural cross-correlation algorithm

For comparative purposes, outputs from a binaural cross-correlation algorithm were calculated (Akeroyd, 2017). The stimuli NoSo, NoSπ, NπSπ, and NπSo were generated for a 35 dB tone and a 60 dB randomly distributed broadband noise. Stimuli were sampled at 20 kHz and were 1 s in duration. Signals were passed through gammatone filters centered at 500 Hz and passed through a model of neural transduction (Meddis et al., 1990). The outputs were then delayed relative to one another, and the cross-products calculated and summated.

### 4.9 Statistical analysis

We performed Student’s two-tailed t-tests (assuming unequal variance) to assess differences between BMLDs. Pearson product-moment correlation was calculated between the average responses of nodes to NoSo, NoSπ, NπSπ, and NπSo, and the delay matched outputs of a binaural cross-correlation algorithm. Correlations were calculated with, and without, local averaging (within 600 μs). Student’s two-tailed t-tests (assuming unequal variance) and two-sample Kolmogorov–Smirnov tests were performed to compare changes in latent node activation values for homophasic/antiphasic stimuli pairs. For the outlined statistical analyses, the criterion for significance was set to p=0.05. Violin plots were used to capture data probability density in Figure 3C and Figure 4S. The lightly shaded underlay in Figure 5A is standard error. In Figure 5F an exponential curve was robustly fit with the least absolute residual method.

## 5 Data Availability Statement

The models and datasets generated and analyzed for this study can be found in our github repository [https://github.com/HearingSciences/BinauralDetection_DNN].

## 6 Conflict of Interest

The authors declare that the research was conducted in the absence of any commercial or financial relationships that could be construed as a potential conflict of interest.

## 7 Author Contributions

Conceptualization, S.S.S. and M.A.A.; Methodology, S.S.S.; Investigation, S.S.S.; Interpretation: S.S.S., J.S. and M.A.A.; Writing – Original Draft, S.S.S.; Writing – Review & Editing, S.S.S., J.S. and M.A.A.; Funding Acquisition, M.A.A.; Resources, M.A.A.; Supervision, M.A.A.

## 8 Funding

S.S.S. and M.A.A. are supported by the Medical Research Council (grant number MR/S002898/1). J.S. is funded by a Nottingham Research Fellowship from the University of Nottingham.

## 9 Acknowledgments

We are grateful for access to the University of Nottingham’s Augusta High Performance Computer service. Thanks to Alan Palmer for assistance in sharing previously published physiology data and to Wilten Nicola for his valuable comments during the writing phase.

## Notes

### Competing Interest Statement

The authors have declared no competing interest.

### Summary of Updates

Addition of a proof of concept model. Revised text and figures.

